# Mutational Landscape of Cancer-Driver Genes Across Human Cancers

**DOI:** 10.1101/2022.09.11.507448

**Authors:** Musalula Sinkala

## Abstract

The cancer driver genes are involved in transforming healthy cells into cancerous cells. The molecular aberrations which lead to cancer involve gain and loss of function mutations in various cancer driver genes. Here, we examine the genome sequences of 20,066 primary tumours representing 43 distinct human cancers to identify and catalogue driver mutations in 729 known cancer genes. We show that the frequency of driver mutations in these genes varies significantly between cancer types. We find that the class of cancer driver genes most frequently mutated are the tumour suppressor genes (94%), followed by oncogenes (93%), transcription factors (72%), kinases (64%), cell surface receptors (63%), and phosphatases (22%). Furthermore, we identify the subset of these genes within which mutations exhibit a co-occurrence or mutually exclusive pattern. Interestingly, we find that patients with tumours with different combinations of driver gene mutation patterns tend to exhibit variable survival outcomes. Here, among the well-studied cancer genes, we showed that patients with tumours with *KRAS* and *TP53* mutations are associated with the worst disease outcomes, and those with *PI3KCA* and *BRAF* mutations are associated with favourable survival outcomes. Besides providing new insights into cancer driver mutations, we unearth mutation patterns associated with disease outcomes and various hallmarks of cancer that bring us closer to fully understanding various forms of cancer.

## Introduction

Cancer is fuelled by mutations in driver genes that control various hallmarks of cancer, including escaping programmed cell death, promoting genome instability and mutations, and proliferative signalling ^1^. Known cancer driver genes (CDGs) include genes that encode cell surface receptors, oncogenes, tumour suppressor genes, kinases, phosphatases, and transcription factors ^2–6^. CDGs of these classes transcribe mRNAs that encode proteins, which function in various oncogenic pathways that fuel oncogenesis by enabling various hallmarks of cancer ^7,8^. These CDG mutations qualitatively or quantitatively alter the function of genes and proteins and, consequently, alter the cellular processes in which these proteins participate.

Over the last few decades, our understanding of the genes, the pathways within which they function, and their role in oncogenesis have grown significantly and, along with this increased interest, concerted efforts to treat various cancer types ^9–14^. Furthermore, various genetic aberrations have been identified in human cancers, and several of the proteins encoded by these genes are well-established drug targets, and others are promising drug targets ^3,4,15,16^. Thus far, while a wide range of known driver mutations is impressive, the list covers only a few cancer types.

Knowledge of driver alterations that impact gene function and the associated consequence would allow us to discover a phenomenological and functional causal chain of events leading to neoplasm. Some of this knowledge has been successfully applied to design therapies specifically targeting proteins altered by somatic and germline mutations in driver genes ^17,18^. However, we still do not completely understand the extent to which driver genes and the classes thereof are altered across the entire spectrum of human cancers and within each cancer type.

Here, we utilise publicly available datasets generated by various cancer sequencing projects to understand the extent of CDGs alterations in human cancers.

Furthermore, we obtain information on known cancer genes compiled by the Catalog of Somatic Mutations in Cancer (COSMIC), Cancer Gene Consensus (CGS) database ^19,20^. Then, we comprehensively analyse known CDG mutations across different cancer types by integrating information on tumour genetic alterations with known gene annotations. Besides providing novel biological insights into the mutational landscape of these cancer genes, we show both the extent to which these are co-occurring or exclusive in tumours of various tissues and their association with the disease outcome of patients.

## Results

### The mutational landscape of cancer genes in human cancers

We first identified a list of known cancer genes based on the information from COSMIC ^19^, and CGC ^21^. The list includes 729 genes that encode oncogenes (377), tumour suppressor genes (TSGs; 350), transcription factors (126), kinases (93), phosphatases (10), and cell surface receptor proteins (CSR; 69) (see Supplementary File 1).

Next, we obtained whole-exome sequencing data from 118 cancer studies focusing on 43 different human cancer types and involving 20,066 samples (Supplementary File 1). The number of samples in each study varied. Among the cancer types, Breast Carcinoma represented the highest number of samples (2,338) and Thymomas the least (102), Figure 1a. We then calculated the somatic mutation frequencies in the 729 genes across the samples of the 43 cancer types (Supplementary File 1) and found that mutation frequency for different cancer genes ranged from 0% for the *A1CF* (and other genes) in Acute Myeloid Leukaemia to 94% for the *TP53* gene in Small Cell Lung Carcinomas (Figure 1b and 1c). Other cancer genes with high mutation frequencies include the *VHL* (mutated in 77%) of Clear Cell Renal Cell Carcinomas and *PTEN* (67%) of Uterine Corpus Endometrial Carcinomas, and *JAK2* (76%) of Myeloproliferative Neoplasms (Figure 1c, also see Supplementary File 1).

**Figure 1:**
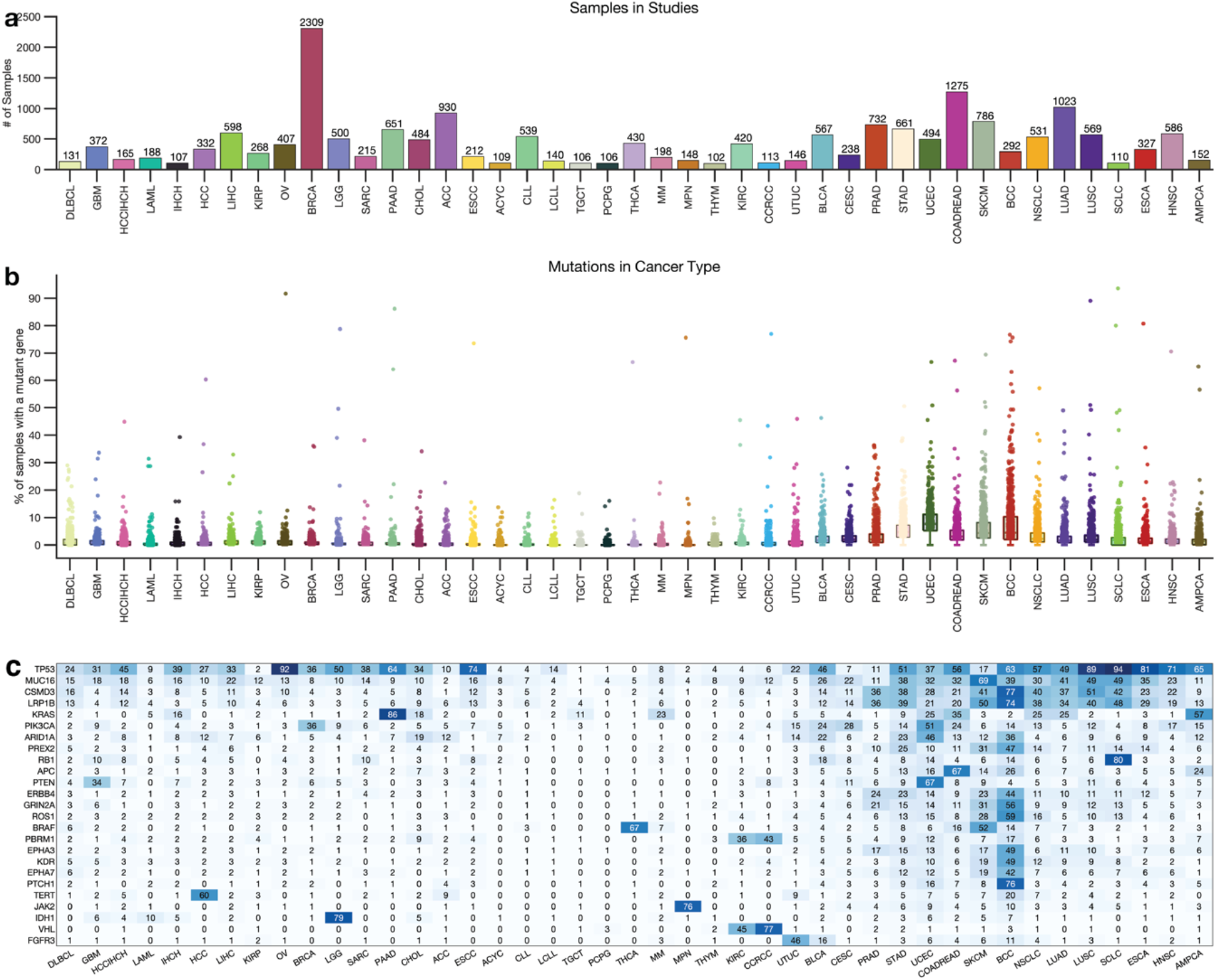
**(a)** Distribution of 20,066 tumours across 43 human cancer types. The disease codes and abbreviations: UCEC, uterine corpus endometrial carcinoma; SKCM, skin cutaneous melanoma; BLCA, bladder urothelial carcinoma; UCS, uterine carcinosarcoma; OV, ovarian serous cystadenocarcinoma; LUSC, lung squamous cell carcinoma; STAD, stomach adenocarcinoma; LUAD, lung adenocarcinoma; ESCA, oesophageal adenocarcinoma; DLBC, diffuse large b-cell lymphoma; CESC, cervical squamous cell carcinoma; HNSC, head and neck squamous cell carcinoma; SARC, sarcoma; LIHC, liver hepatocellular carcinoma; BRCA, breast invasive carcinoma; COADREAD, colorectal adenocarcinoma; CHOL, cholangiocarcinoma; ACC, adrenocortical carcinoma; PAAD, pancreatic adenocarcinoma; PRAD, prostate adenocarcinoma; GBM, glioblastoma multiforme; KIRP, kidney renal papillary cell carcinoma; KIRC, kidney renal clear cell carcinoma; MESO, mesothelioma; LGG, brain lower grade glioma; UVM, uveal melanoma; PCPG, pheochromocytoma and paraganglioma; TGCT, testicular germ cell tumours; KICH, kidney chromophobe; THYM, thymoma; LAML, acute myeloid leukaemia; THCA, thyroid carcinoma. **(b)** Mutation frequency of known cancer driver genes within each human cancer type. **(c)** Clustered heatmap of the most frequency mutated cancer genes across types using the percentage of tumours with genes displaying alterations. Increasing colour intensities denote higher percentages. The heat map was produced using unsupervised hierarchical clustering with the Cosine distance metric and complete linkage (see Figure S1).

Next, for each cancer type, we summarised the number of mutated genes in 1) none of the samples, 2) less than 5 per cent of the samples, and 3) more than 5% of the samples. Interestingly, we found that most driver cancer genes are not mutated in some cancer types, and fewer than 10 genes are mutated in more than 5% of the samples. For example, only two known cancer genes in Thymomas (*MUC16* and *HRAS*), Testicular Germ Cell Tumours (*KRAS* and *KIT*), and Thyroid Carcinomas (*BRAF* and *NRAS*) were mutated in more than 5% of the tumours (Figure 1c, Supplementary Figure 1, and Supplementary File 1).

Furthermore, some cancer types had a significantly higher number of known cancer genes mutated in more than 5% of the samples, e.g., in Uterine Corpus Endometrial Carcinoma (568 known cancer genes mutated), Stomach Adenocarcinoma (330 genes), Skin Cutaneous Melanoma (321 genes), and Basal Cell Carcinoma (344 genes). This finding shows that the extent to which the cancer genes are mutated across cancer types varies and that some cancer types have few mutations within the coding sequences of known cancer genes ^22,23^.

### Mutations of each cancer gene across all tumours

We calculated the mutation frequency of each cancer gene across 43 different human cancer types in all the 20,066 samples. We found that 98.9% (719 out of 726) of cancer genes were mutated in at least one sample. Among the oncogenes, *MUC16* (mutated in 20.9% of tumours), *PIK3CA* (13.0%), and *KRAS* (10.7%) were the most frequently mutated across the profiled samples (Figure 2a). Furthermore, among the TSGs, *TP53* (38.2%), *CSMD3* (17.2%), and *LRP1B* (16.9%) were the most frequently mutated (Figure 2b). *PIK3CA* (13.0%), *BRAF* (7.3%), and *ERBB4* (7.2%) were the top-three mutated genes that encode protein kinases (Figure 2c).

**Figure 2:**
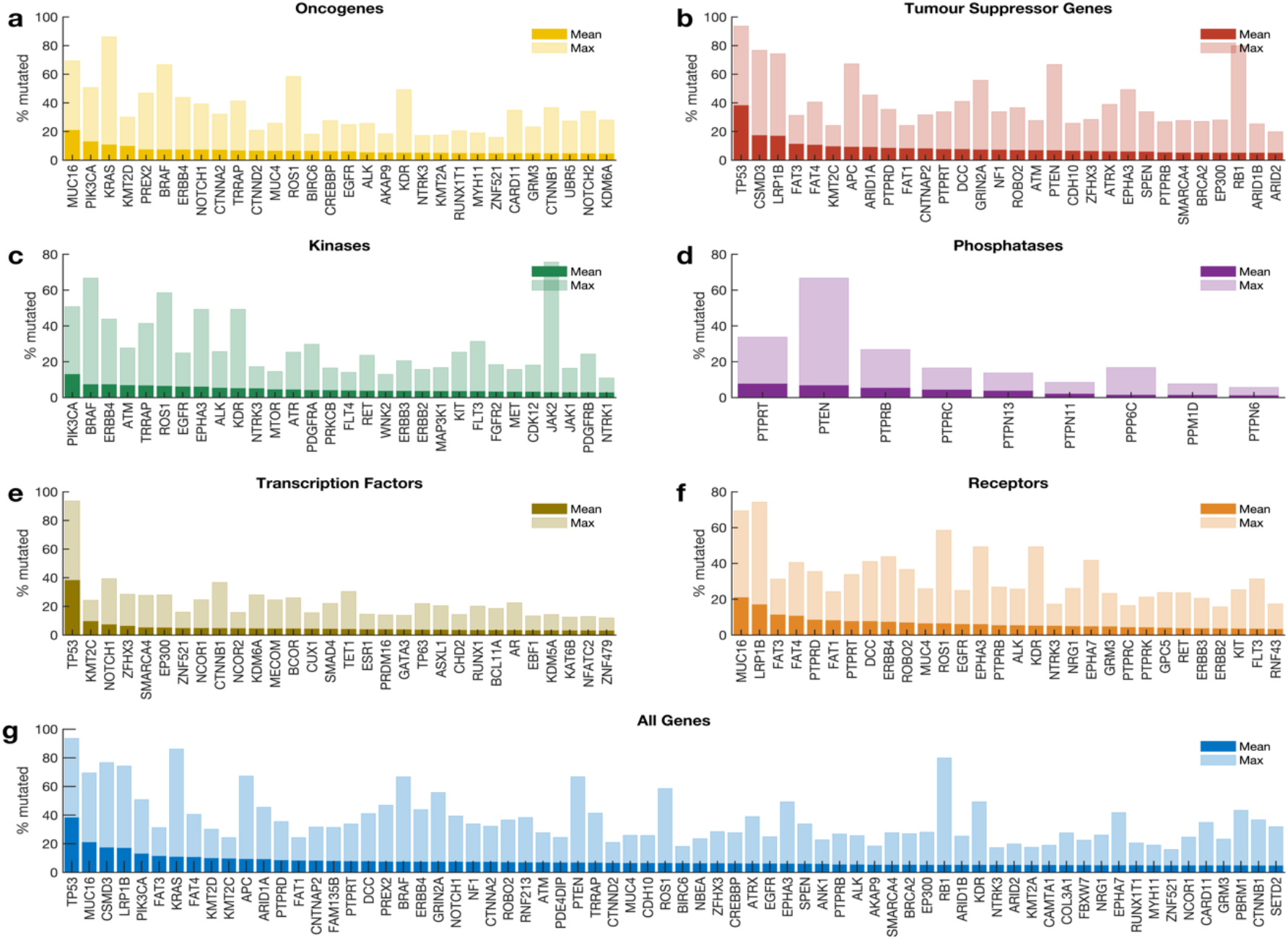
Frequency of known driver gene mutations among: **(a)** oncogenes, **(b)** tumour suppressor genes, genes that encode **(c)** kinases, **(d)** phosphatases, **(e)** transcription factors, **(f)** cell surface receptor proteins, **(g)** all the genes.

Furthermore, we found that *PTPRT* (7.6%) and *PTEN* (6.7%) were the most frequently mutated among the genes that encode protein phosphatases (Figure 2d), whereas *TP53* (38.2%) and *NOTCH1* (7.2%) were the top-two frequently mutated among the genes that encode transcription factors (Figure 2e). Also, among the genes that encode cell surface receptors, we found that *MUC16* (20.9%) and *LRP1B* (16.9%) were the most frequently mutated (Figure 2f). Overall, the top-five frequently mutated cancer genes across human cancers were *TP53* (38.2%), *MUC16* (20.9%), *CSMD3* (17.2%), *LRP1B* (16.9%), and *PIK3CA* (13.0%) (Figure 2g).

The mutation frequencies we report here are reasonably consistent with previous reports, which indicated that *TP53* (34.5% across all samples) is the most frequently altered gene, followed by *PIK3CA* (13.0%) ^3,24^. Furthermore, we found that the extent to which the cancer genes are mutated in different cancer types varies significantly, a pattern likely to impact the treatment strategies that could be applied to cancer of different tissues ^25^.

### Mutations in categories of cancer genes

We were interested in evaluating the extent to which genes in particular categories of CDGs (oncogenes, TSGs, transcription factors, kinases, phosphatases, and receptors) are mutated across human cancers. Here, we found mutations in the known cancer genes in all tumours. Of the six classes of cancer genes, the TSGs (94% of the tumours) and the oncogenes (93%) showed the highest frequency of mutations, followed by the transcription factors (72%), kinases (64%), receptors (63%), and the phosphatases (22%); (Figure S2 and Figure 3a). Overall, our analyses revealed that the mutational landscape of the six cancer gene classes was mainly consistent within cancer (Figure 3a). Therefore, we suggest that the observed correlation in mutation frequencies between CDGs of different classes in a particular cancer type may indicate that gene mutations tend to co-occur.

**Figure 3:**
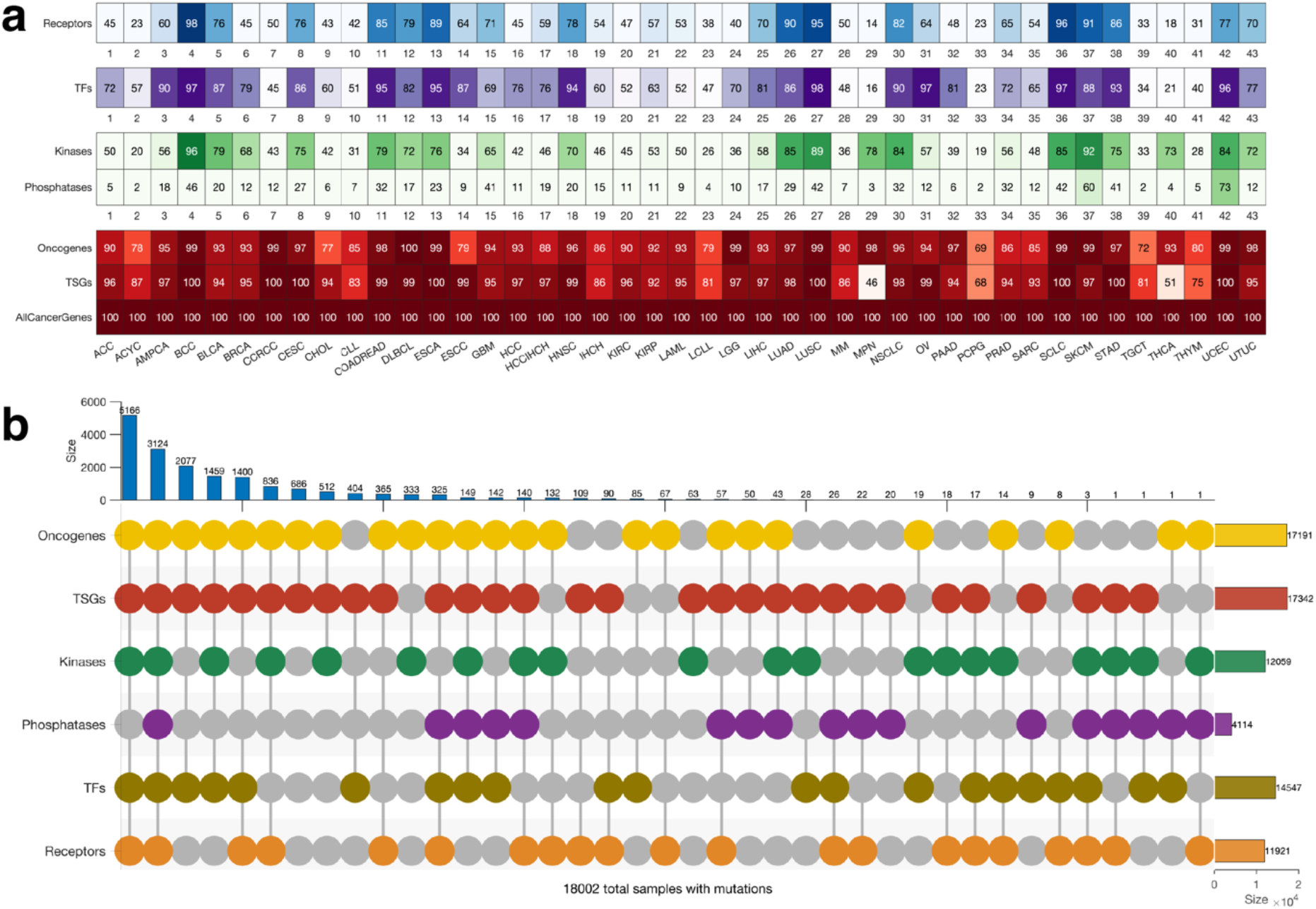
**(a)** Frequency of tumours of different cancer types with altered genes that encode (from top to bottom) cell surface receptors, transcription factors, kinases, phosphatases, oncogenes, tumour suppressor genes and all the cancer driver genes. **(b)** Upset plot showing the mutual exclusivity and co-occurrence of mutations across the different classes of cancer driver genes. Note that the counts comprise only mutations in tumours with mutations in genes involved in more than one of the six cancer gene classes. See Figure S2 for the number of mutations exclusive to each class of genes.

Also, we found that 96% (18,644 samples) of all the tumours harboured mutations in genes involved in more than one of the six cancer gene classes (Figure 3b). Furthermore, we found 3,085 tumours harbour mutations in all six classes of genes, and another 4,978 tumours harbour mutations in all six classes of genes, except those that encode protein phosphatases (Figure 3b). Conversely, 778 samples harboured mutations in only one class of the known cancer genes (Figure S3). Overall, our finding shows that for most cancer types, the tumours tend to have mutations in genes of at least five of the six classes of the CDGs.

### Co-occurrence and exclusivity of mutations in cancer gene pairs

Given that we found a convolved pattern in the mutational landscape of the known cancer genes (Figure 4a and Supplementary Figure 3a and 3b), we were interested in determining the extent to which gene mutations tend to be mutually exclusive or co-occurring. First, we evaluated the pattern of mutations for pairs of genes across the 43 different human cancer types in all the 20,066 samples (see methods sections). Here, we found mutations in 150,728 gene pairs significantly co-occurring, 21 mutually exclusive gene pairs, and 226 gene pair mutations that were non-statistically significant. Among the significantly mutually exclusively mutated gene pairs were *BRAF* and *TP53* (p = 2.4 × 10^−12^), *BRAF* and *KRAS* (p = 1.5 × 10^−7^), and *TP53* and *NRAS* (p = 1.5 × 10^−05^), also see Supplementary File 2.

**Figure 4:**
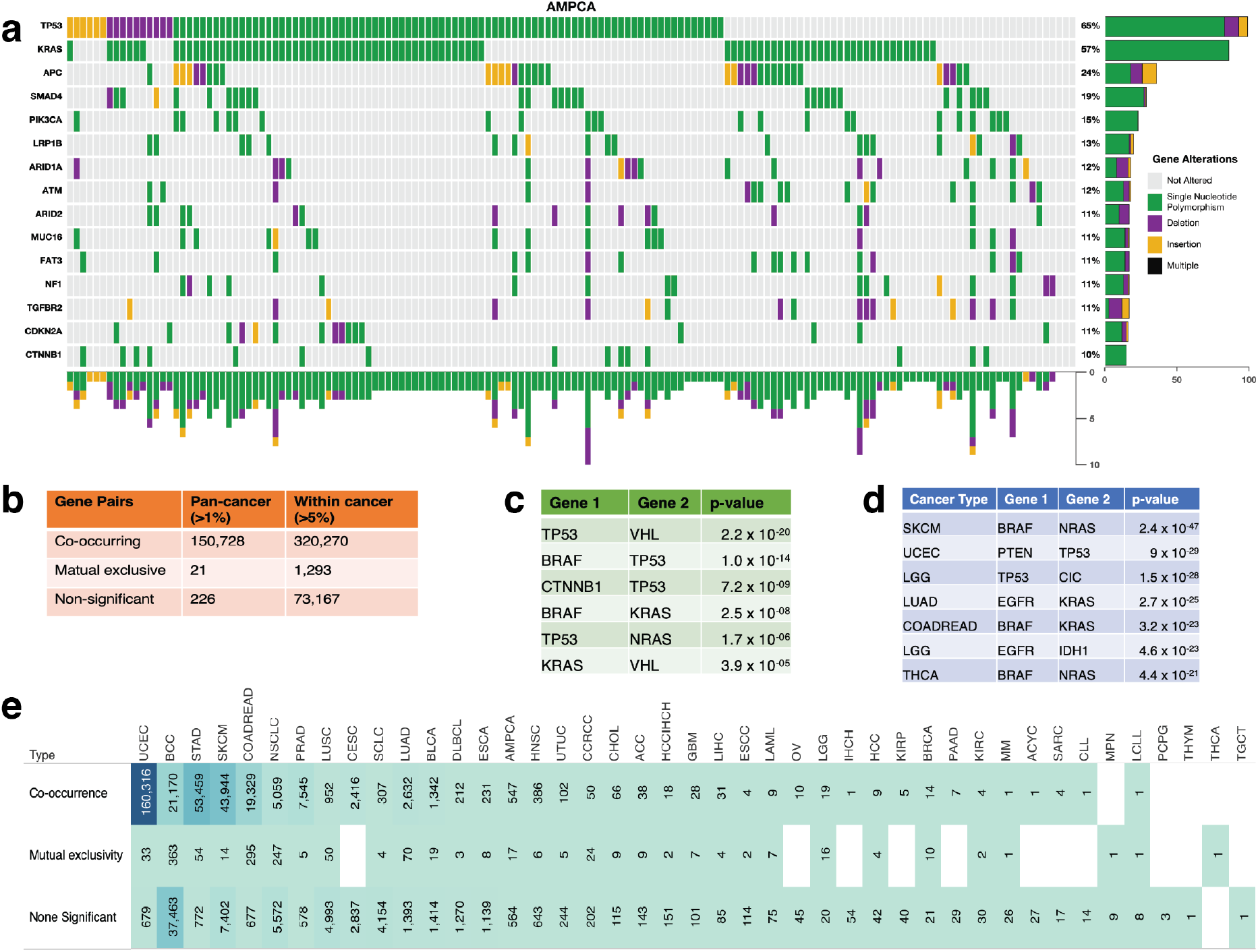
**(a)** Example of mutation signature plot showing co-occurrence and exclusivity of mutations in ampullary carcinoma showing the 13 most frequently mutated genes. Shown along the plot column are samples that have mutations along the rows. Also, see supplementary Figure 4a and 4b. **(b)** The number of identified co-occurring and mutually exclusive mutations across all cancer types and within each cancer. **(c)** Top-7 most significantly co-mutated gene pairs across Pancancer studies. **(d)** Top-7 most significantly mutually exclusively mutated gene pairs within each cancer study. **(e)** The number of driver gene mutations that are statistically significantly co-occurring, mutually exclusive, and not significantly co-occurring or mutually exclusive (none).

Next, for each of the 43 cancer types, we assessed the exclusivity or co-occurrence of mutations in gene pairs we found mutated in more than 5% of the tumours. We found 1,523 gene pairs with significantly co-occurring mutations, 213 gene pairs with significantly mutually exclusive mutations, and 4,206 gene pairs that exhibit a non-statistically significant mutation pattern (Figure 4a). Among the top three exclusively mutated gene pairs are *BRAF* and *NRAS* in Skin Cutaneous Melanoma (p = 2.4 × 10^−47^), *PTEN* and *TP53* in Uterine Corpus Endometrial Carcinoma (p = 9.0 × 10^−29^), and *TP53* and *CIC* in Brain Lower Grade Glioma (p = 1.5 × 10^−28^). Furthermore, we found that in particular cancer types, some gene pairs exhibit a significantly co-occurring mutation pattern, including *TP53* and *ATRX* in Brain Lower Grade Glioma (p = 4 × 10^−57^), *TP53* and *CDKN2A* in Head and Neck Squamous Cell Carcinoma (p = 1.3 × 10^−11^), *KRAS* and *PIK3CA* in Colorectal Adenocarcinoma (p = 4.2 × 10^−06^) and Uterine Corpus Endometrial Carcinoma (p = 2.9 × 10^−05^). See Supplementary File 2 for the list of mutually exclusive and co-occurring mutated gene pairs in specific cancer types.

Overall, the higher number of exclusive driver mutations we found within cancer types (213 gene pairs) compared to across all cancer types (8 gene pairs) may indicate a selection for mutations in specific driver gene pairs in specific cancer types 26. Furthermore, we suggest that the exclusively mutated gene pairs dysregulate divergent oncogenic pathways in specific cancer types ^27,28^.

Next, we calculated the number of co-occurring and exclusively mutated gene pairs within each cancer type. Here we found that Basal Cell Carcinoma (363 gene pairs) has the highest number of exclusively mutated gene pairs among the 43 cancer types, followed by Colorectal Adenocarcinoma (295 pairs) and Non-Small Cell Lung Cancer (247 pairs). Conversely, Uterine Corpus Endometrial Carcinoma (160,316 pairs) displayed the highest number of co-occurring mutations in gene pairs, followed by Stomach Adenocarcinoma (53,459 pairs) and Skin Cutaneous Melanoma (43,944 pairs), see Supplementary File 2.

Interestingly, we found that some gene pairs may exhibit an exclusive mutation pattern in one cancer type and a co-occurrence pattern in another. For instance, we found that mutations of *TP53* and *PIK3CA* tended towards mutual exclusivity in Breast Carcinoma, Colorectal Adenocarcinoma, and Brain Lower Grade Glioma but co-occurred in Non-Small Cell Lung Cancer (Figure 5a). Furthermore, *TP53* and *KRAS* mutations co-occur in Lung Adenocarcinoma and Pancreatic Ductal Adenocarcinoma but are mutually exclusive in Uterine Corpus Endometrial Carcinoma and Cholangiocarcinoma (Supplementary File 2). Here, we suggest that in cases where the gene pairs are co-mutated, they may function together to promote oncogenesis ^29–32^. Conversely, when exclusively mutated, the gene pairs may act independently to promote oncogenesis, yielding tumours of different phenotypic or molecular subtypes in a particular tissue ^27,28,33^.

**Figure 5:**
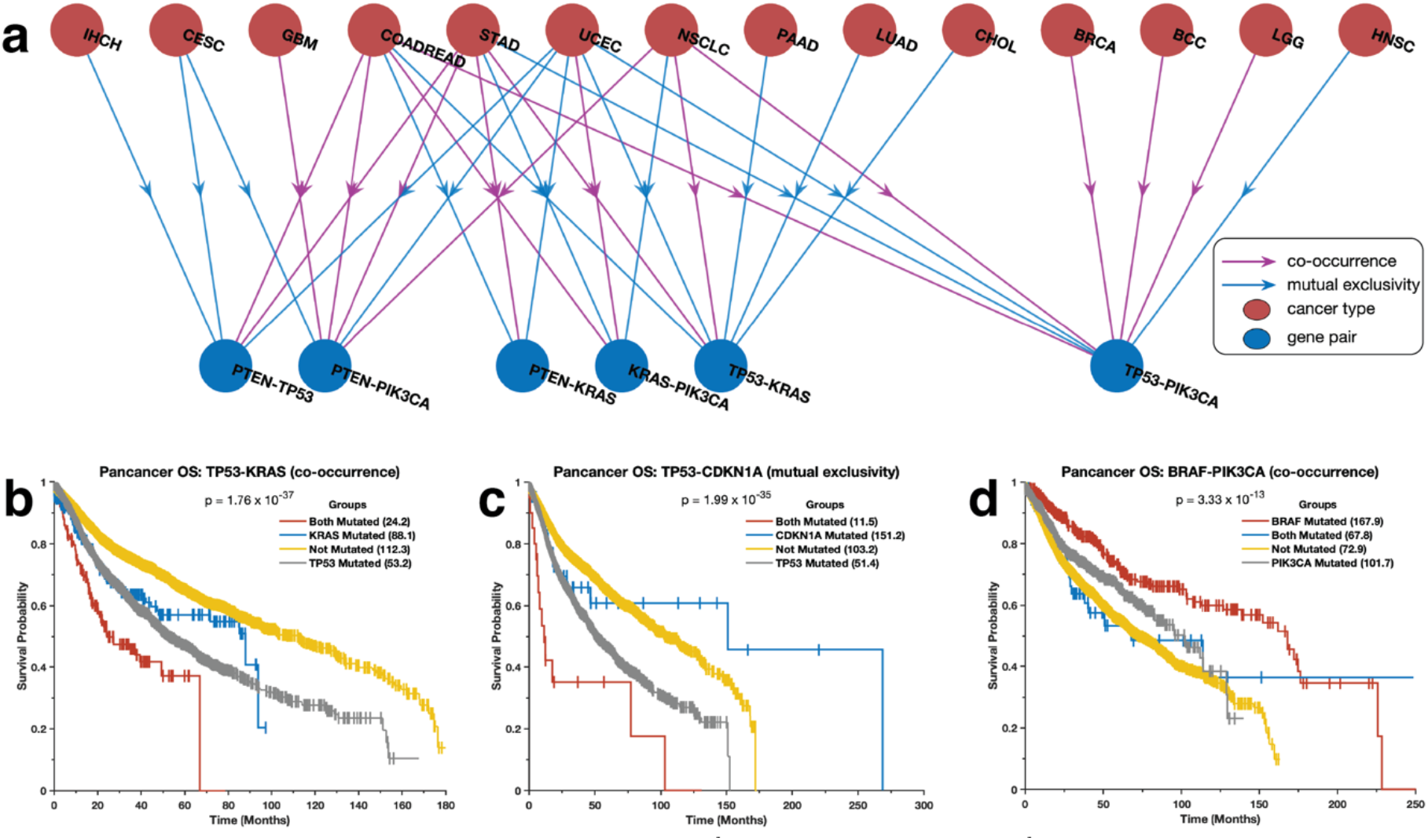
**(a)** The co-occurrence and exclusivity of mutations in some of the most studied driver gene pairs in different cancer types. Kaplan-Meier curve of the disease-free survival periods and overall survival periods of patients afflicted by tumours that: **(a)** both have TP53 and KRAS mutations, only TP53 mutation, only KRAS mutations, and no KRAS or TP53 mutations; **(b)** both have PI3KCA, and EGFR mutations, only PI3KCA mutation, only EGFR mutations, and no PI3KCA or EGFR mutations; **(c)** both have PI3KCA, and EGFR mutations, only PI3KCA mutation, only EGFR mutations, and no PI3KCA or EGFR mutations; **(d)** both have PI3KCA and EGFR mutations, only PI3KCA mutation, only EGFR mutations, and no PI3KCA or EGFR mutations.

Additionally, we supposed that the number of exclusive mutations should correlate with co-occurring mutations across the cancer types. However, this is not the case (see Supplementary Figure 5a). Furthermore, there was no apparent correlation between the number of cancer-type samples with the number of exclusively mutated gene pairs we observed (Supplementary Figure 5b). This absence of correlation is pronounced by the mutation pattern in Uterine Corpus Endometrial Carcinoma, where we found 160,316 co-occurring gene pairs and only 33 exclusive gene pairs across 661 samples. Our results suggest that for a particular form of cancer, the extent of mutually exclusively mutated gene pairs indicates the genomic complexity of the disease linked to alterations in different oncogenic pathways.

### Mutation patterns are associated with disease outcomes

We assessed the impact of mutations in gene pairs on the overall survival of cancer patients (see Methods section). Briefly, we grouped the patients into four groups based on the mutations in a gene pair: 1) no mutations, 2) and 3) only one of the genes in the pair is mutated, and 4) both genes are mutated. Interestingly, we found that mutations in gene pairs are associated with varied overall survival durations of patients afflicted. For example, we found that patients with tumours that harbour mutations in both *KRAS* and *TP53* (p = 1.76 × 10^−37^) or *CDKN1A* and *TP53* (p = 1.99 × 10^−35^) tended to exhibit worse survival outcomes than those with tumours in which one or none of these genes are mutated (Figure 5b and 5c, see Supplementary File 3). Furthermore, we found that patients with tumours with mutations in *PIK3CA* and/or *BRAF* tended to exhibit better survival outcomes than those with mutations in *TP53, KRAS*, and/or *EGFR* (Figure 5c and Supplementary Figure 6). In addition, we found that the patients with tumours with mutations in *PIK3CA, BRAF, CDH1*, and *NRAS* exhibit better survival outcomes than those without mutations in these genes (Supplementary Figure 6; and Supplementary File 3). Our findings show that some cancer patients may exhibit significantly favourable disease outcomes due to their tumours’ mutation profile.

### Driver pathways of co-occurring and exclusively mutated genes

Given that we found the co-occurrence and exclusivity of mutations in gene pairs intriguing, we identified the driver pathways (gene combinations) for the top 10 most frequently mutated CDGs in each cancer type (see Methods sections and Supplementary File 4). For example, we found two gene sets of co-occurring mutated driver pathways in Ampullary Carcinoma (Figure 6a). The first of these pathways involves five genes (*TP53, KRAS, APC, SMAD4*, and *PIK3CA*) exhibiting a co-occurring mutation pattern and the second set of five genes (*ARID1A, ATM, ARID3, NF1*, and *TGFBR2*) that are exclusively mutated. In addition, in Acute Myeloid Leukaemia, we found two driver gene sets; the first set includes six genes (*FLT3, DNMT3A, NPM1, IDH3, RUNX1*, and *IDH1*) that exhibit a co-occurring mutation pattern, and the second set of four genes (*TET2, TP53, NRAS*, and *WT1*), exhibit an exclusive mutation pattern (Figure 6b). Furthermore, in Lung Adenocarcinoma, we found two driver gene sets; the first set includes six genes (*TP53, EGFR, KRAS, KEAP1, STK11*, and *NF1*) that exhibit a co-occurring mutation pattern, and the second set of four genes (*SMARCA4, ATM, RBM10*, and *APC*) (Figure 6c), exhibit an exclusive mutation pattern.

**Figure 6:**
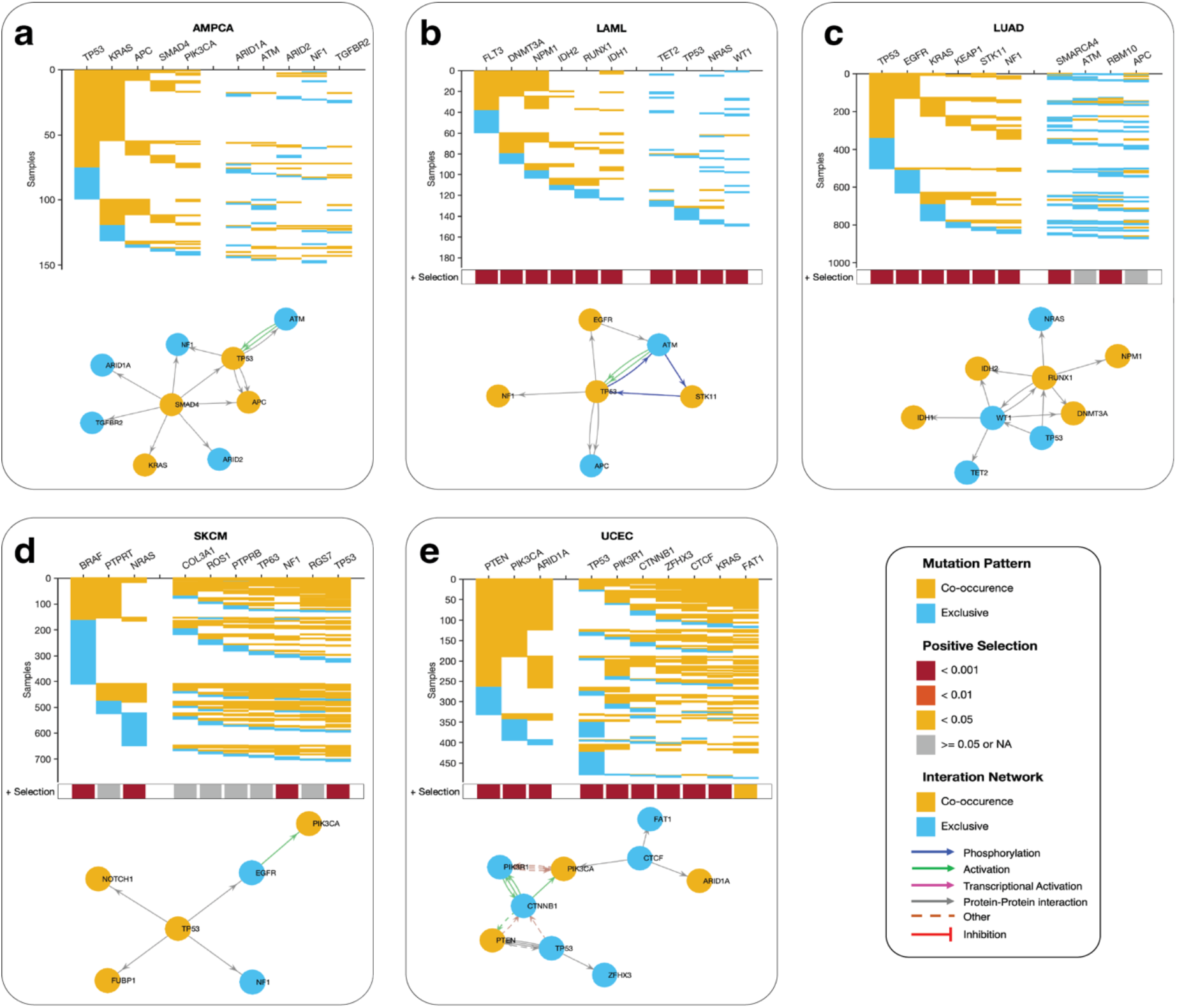
The two co-occurring pathways that are predicted to drive oncogenesis based on the mutational landscape in **(a)** AMPCA, **(b)** LAML, **(c)** LUAD, **(d)** SKCM, and **(e)** ECEC. The coloured square marks at the bottom of each plot show a positive selection of mutations in each gene along each column (see Methods section). The connectivity of network components within each panel was extracted from the KEA and ChEA databases and the UCSC super pathway.

However, we found that driver gene mutations tend to significantly co-occur in some cancer types, including Uterine Corpus Endometrial Carcinoma, Skin Cutaneous Melanoma, and Basal Cell Carcinoma (Figure 6d and 6e, also see Supplementary Figure 7). Interestingly, we found that mutated genes of cancer driver pathways are significantly under positive selection across all cancer types.

### Driver gene mutations in the context of cancer hallmarks

The hallmarks of cancer are biological abilities acquired during tumour development that impose a substantial degree of complexity on cancer. Owing to the importance of cancer hallmarks in efforts to design better treatment strategies, we set to define the extent to which genes of each hallmark of cancer are altered across human cancers (Figure 7 and Supplementary File 5, see Methods section). Here, we found that the highest number of mutated genes are in the hallmark “escaping programmed cell death” genes (213 genes), followed by “invasion and metastasis” (204 genes), proliferative signalling (160 genes), and “genome instability and mutations” (121 genes), see Figure 7. Interestingly, among these cancer hallmarks, the most frequently mutated genes are oncogenes and tumour suppressor genes that are not kinases, phosphatases, or cell surface receptors. We find this interesting because most effectors in cancer research to identify drug targets focus on kinases and cell surface receptors. Our findings demonstrate the potential to identify many variable drugs among the non-traditionally targeted CDGs.

**Figure 7:**
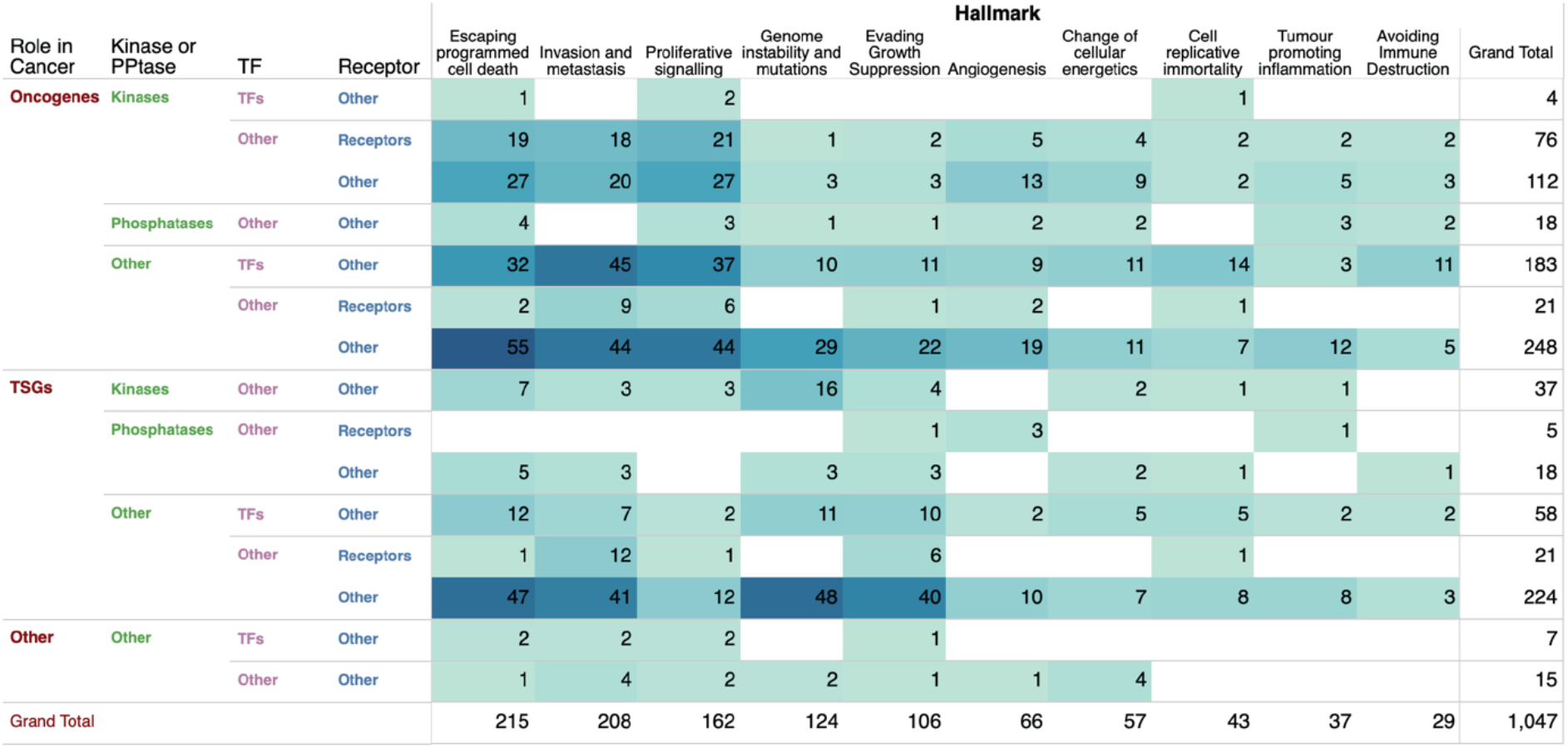
The total number of cancer driver genes across each combination of cancer genes classes associated with the hallmarks of cancer.

## Discussion

We have conducted a systematic analysis of the 729 CDG mutations across 43 human cancers. Here, we found at least one non-synonymous mutation in known CDGs in all analysed samples. Furthermore, we found mutations in oncogenes, TSGs, gene encoding transcription factors, kinases, phosphatases, and cell surface receptors in all the cancer types we analysed. This finding demonstrates that various components of the cell signalling processes, including receptors that respond to stimuli, cytoplasmic enzymes, and nuclear proteins, are involved in oncogenesis. Interestingly, we found that not all samples of a particular cancer type harbour the same driver mutations, and the distribution of gene mutations within each cancer type varies significantly. With this finding, we suggest that each patient may exhibit a different combination of mutations sufficient to perturb various oncogenic pathways, and therefore, understanding the mutation profile of each patient’s tumour should be crucial for optimising personalised cancer treatments.

Our findings emphasize the significance of the CDG mutations across human cancers. However, it must be stressed that some tumours have less than 5% of CDGs mutated. For example, gene mutations are infrequent in Thyroid carcinoma, Testicular Germ Cell Tumours, and Thymomas, where only two driver genes are mutated in more than 5% of the examined tumours. These exceptions reinforce the notion that multiple routes to oncogenesis may be independent of cancer gene mutations, and involve alterations in other regulatory mechanisms such as the epigenome ^1,34–36^. Furthermore, these findings reveal that, in the case where driver gene mutations are infrequent, we need to look for CDGs using other genetic datasets, e.g., DNA methylation profile and mRNA transcription abundance and/or mutations in the non-coding and regulatory regions of the genome.

It should be noted that despite the large amounts of genomic data we analyse here, we could not pinpoint the commonly mutated driver genes in samples of specific cancer types. Therefore, we are impeded from identifying commonly applicable drug targets and marker mutations within each cancer type because of the sparsity nature of gene mutations and the limited diversity of the presently available genome sequences.

We further showed that co-occurrence and exclusive nature of driver mutations significantly affect the disease outcome of the afflicted patients for various forms of cancer. These findings show that various gene alterations of specific gene pairs have a diverse impact on processes that drive disease aggressiveness.

Altogether, there is a pressing need for new molecular tools that would allow us to describe the impact of different combinations of gene alterations on disease aggressiveness and the chemosensitivity of tumours. Despite the currently available large genomics data, we still cannot study the impact of every combination of gene mutations because such information is unavailable. Future experiments that would allow us to alter cancer genes in normal cells in different combinations would eventually allow us to unlock the impact that a combination of driver mutations has on oncogenesis, disease aggressiveness and the chemosensitivity of tumours.

## Methods

We analysed 20,066 patient-derived tumour samples representing 43 distinct human cancers, obtained from cBioPortal ^37^ version 3.1.9 (http://www.cbioportal.org; see Supplementary File 1 for details on the cancer studies). The elements of the data that we obtained from cBioPortal include somatic gene mutations (point mutations and small insertions/deletions) and comprehensively deidentified clinical data.

### Compilation of cancer-driver genes and their classes

We obtained information on cancer-driving genes from the Catalogue of Somatic Mutations in Cancer^19^ (COSMIC) Cancer Gene Census (CGC)^21^. The CGC datasets comprised 729 cancer-driving genes that are evidence-based and manually curated (COSMIC v95, released 24-NOV-21). Next, we set to classify these known cancer genes into five classes, i.e., oncogenes, tumour suppressor genes, kinases, phosphatases, cell surface proteins, and transcription factors. Therefore, we obtained information of the gene annotations from the various databases, including the (1) the Sanger Consensus Cancer Gene Database^19^ (699 oncogenes and TSGs) ; (2) the UniProt Knowledgebase^38^ (304 oncogenes and 741 TSGs); (3) the TSGene database ^39^ (1,220 TSGs); (4) ChEA transcription factor database ^40^ (645 transcription factors); TF2DNA database ^41^ (1,314 transcription factors); (5) Kinase Enrichment Analysis database ^42^ (428 kinases); (6) the ONGene database^43^ (725 oncogenes); and Surfaceome database ^44^ (2,950 cell surface receptors). Then, we collated the driver genes from all the above databases to obtain a list of xx driver cancer genes (after removing overlapping genes), including 384 oncogenes, 370 tumour suppressor genes, 10 phosphatases, 93 kinases, 73 cell surface receptors, and 203 transcription factors (Supplementary File 1).

### Calculation of cancer gene mutations

We obtained whole-exome sequencing of the samples for all the cancer driver genes. Next, we returned the non-synonymous mutations that occurred within the genes. Then, to evaluate the extent to which each cancer drive gene is mutated in cancer, we calculated the somatic mutation frequency for each gene across the 20,066 samples across each cancer type (Supplementary File 1). Furthermore, for each cancer type, we obtained summaries of the number of mutated genes in 1) none of the samples, 2) less than 5 per cent of the samples, and 3) more than 5% of the samples.

### Cancer driver gene mutations across gene classes

We sort to determine the extent to which the cancer driver genes were mutated within each cancer type and across all human cancer. Here, we calculated the frequency of non-synonymous somatic mutations for genes categories such as oncogenes, tumours suppressor genes, kinases, phosphatases, cell surface receptors, and transcription factors. (Supplementary File 1).

### Assessment of the co-occurrence and exclusivity of gene mutations

As described elsewhere, we used the hypergeometric Fisher test to evaluate the correlation in the mutation profile of cancer gene pairs. First, we obtained a list of mutated genes in more than 1% (550 cancer driver genes) of all tumours across all the samples. Then, we applied the Fisher test to each pair of the selected genes and utilised a cut-off p-value of 0.05 to identify statistically significant gene pair correlations. Furthermore, we used the magnitude of the odds ratio to identify gene pairs with co-occurring mutations (odds > 1 and p < 0.05) and gene pairs with mutually exclusively occurring mutations (odds < 1 and p < 0.05). Also, we used the approach to identify driver cancer gene pairs within co-occurring or mutually exclusive mutation patterns within each of the 43 human cancer types (see Supplementary File 2).

### Correlation between mutations pattern and disease outcomes

The Kaplan-Meier method was used to estimate the duration of the overall survival and disease-free survival of patients with cancer with 1) only one gene pair mutated, 2) the other gene pair mutated, 3) none of the gene pair mutated, and 4) both gene pairs mutated.

### Identification of exclusive and co-occurring driver pathways

To discover cancer driver gene combinations based on the patterns of mutations associated with each cancer type, we applied the CoMDP algorithm, which employs a mathematical programming method to identify *de novo* driver pathways in cancer from mutation profiles ^45^. Briefly, the method identifies pathways with mutated cancer driver genes with both high coverage (i.e., present in multiple samples) and high exclusivity, and the pathways exhibit a statistically significant co-occurrence pattern. We used the top-10 mutated driver genes as input for CoMDP algorithms for each cancer type. These top-10 genes were selected from the significantly mutated genes identified using the MutSigCV2 algorithm. In cases where MutSigCV2 identified N genes significantly mutated genes, where N is less than 10, we included N-10, the other most frequently mutated genes, to upsize the gene set size to 10. If not identified to be significantly mutated using MutSigCV2, the following genes were excluded from the analysis: *CSMD1, CSMD3, NRXN1, NRXN4, CNTNAP2, CNTNAP4, CNTNAP5, CNTN5, PARK2, LRP1B, PCLO, MUC16, MUC4, KMT2C, KMT2A, KMT2D, FAT1, FAT2, FAT3*, and *FAT4*. Then, we ran the CoMDP test for each cancer type with *K* = 10, where *K* equals the gene set size. Here, the CoMDP analysis returned mutated driver pathways associated with the genes in each cancer type (Supplementary File 4). Furthermore, information on the selectivity of gene mutations in each cancer type was obtained from the supplementary data of Martincorena et al. ^26^.

### Association between cancer gene mutations and the cancer hallmarks

We accessed information on genes and proteins of various Hallmarks of Cancer from the Catalog of Somatic Mutations in Cancer (COSMIC) database^19^. Then, we calculated the number of mutated genes within each class based on the various cancer hallmark gene sets. Furthermore, we calculated instances where mutations in genes of multiple cancer hallmarks co-occur or are mutually exclusively mutated in human tumours (Supplementary File 5).

### Statistics and Reproducibility

We performed all statistical analyses in MATLAB 2022a. To indicate statistical significance, all comparisons were made using two-sided statistical tests with a p-value < 0.05. In addition, we corrected for multiple statistical testing using a two-sided q-value for each comparison using the Benjamini & Hochberg procedure.

## Supporting information

Description of Supplementary Files

## Data Availability

The data that support our results are available in this manuscript, the supplementary data, and from the following repositories: cBioPortal; https://www.cbioportal.org/, and the COSMIC Consensus Cancer Genes; https://cancer.sanger.ac.uk/census.

## Code Availability

Custom code written in MATLAB for processing and analysing the presented data is freely available at https://github.com/smsinks/Analysis-of-Cancer-Gene-Mutations. In addition, the repository includes some pre-downloaded datasets and conversion files required for the analysis.

## Ethics approval

The study protocol was approved by The University of Cape Town; Health Sciences Research Ethics Committee IRB00001938. The publicly available datasets were collected by the cBioPortal, and TCGA projects and made available via their respective project databases. The methods used here were performed following the relevant policies, regulations and guidelines provided by the TCGA, CCLE, DepMap, GDSC, and LINCS projects.

## Acknowledgements

Funding for this project was provided by H3ABioNet, supported by the National Institutes of Health Common Fund under grant number U24HG006941. The content of this publication is solely the authors’ responsibility and does not necessarily represent the official views of the National Institutes of Health.

