## Supplementary material for "Mutational Landscape of Cancer-Driver Genes Across Human Cancers": Description of Supplementary Files

**Description of Additional Supplementary Files**

**Supplementary File 1:** Supplementary data of cancer studies and mutations of the MAPK pathway genes. The spreadsheet contains the following results/datasets according to the sheet name. ***Cancer Genes:*** list of known cancer driver genes obtained from the COSMIC database[1]: the official HUGO gene symbol, and the class of protein encoded by the genes. ***Cancer studies:*** list and description of individual cancer studies from which our analyses are based. ***Within Cancer Mutations;*** the frequency of mutation and percentage of tumours that harboured mutations in each known cancer gene. ***OGs and TSGs***; mutations that we found in OGs and TSGs across all the cancer studies. **Kinases and Phosphatases**; mutations found in **kinases and phosphatases** across all the cancer studies. **Transcription Factors**: mutations found in **transcription factors** across all the cancer studies. ***Each Gene Mutation***; overall mutations frequencies across all cancer types for each gene. ***Mutation freq grouped;*** frequency of gene mutations across cancer types related to Supplementary Figure 1.

**Supplementary File 2:** The spreadsheet contains the following results/datasets according to the sheet name. ***Each Gene Pair in Cancer Type;*** Mutation pattern of each gene pair across each cancer type. ***Summary of Mutation Pattern;*** **the** number of mutually exclusive, co-occurring, and non-statistically significantly mutated genes across each cancer type.

**Supplementary File 3:** Clinical outcomes across various groups: The spreadsheet contains the following results/datasets according to the sheet name. ***PanCancer OS;*** Pancancer overall survival for patients with tumours categorised into four groups based on the mutations in a gene pair: 1) no mutations, 2) and 3) only one of the genes in the pair is mutated, and 4) both genes are mutated. ***panCancer MultipCompare OS;*** Pairwise comparisons calculated using the Log-rank test [2] for the duration of overall survival periods between patients grouped as described above and related Figure 5 and Supplementary Figure 6. ***PanCancer DSF;*** Pancancer disease-free survival for patients with tumours categorised into four groups based on the mutations in a gene pair: 1) no mutations, 2) and 3) only one of the genes in the pair is mutated, and 4) both genes are mutated. ***PanCancer MultiCompare DFS;*** Pairwise comparisons calculated using the Log-rank test [2] for the duration of disease-free survival periods between patients grouped as described above.

**Supplementary File 4:** Co-occurring gene sets mutated in each cancer type. Gene sets 1 and 2, represent co-occurring driver pathways in each cancer type. n1 and n2 denote the coverage of genomic alterations in each cancer type, and $r_{1,2}$ is the ratio of the common coverage to their union coverage (i.e., co-occurrence ratio). Co-occurrence P represents the p-value of the co-occurrence significance of both pathways

**Supplementary File 5:** Cancer driver gene mutations across various cancer hallmarks. It is related to Figure 7.
