## Supplemental Information for "Mutational Landscape of Cancer-Driver Genes Across Human Cancers"

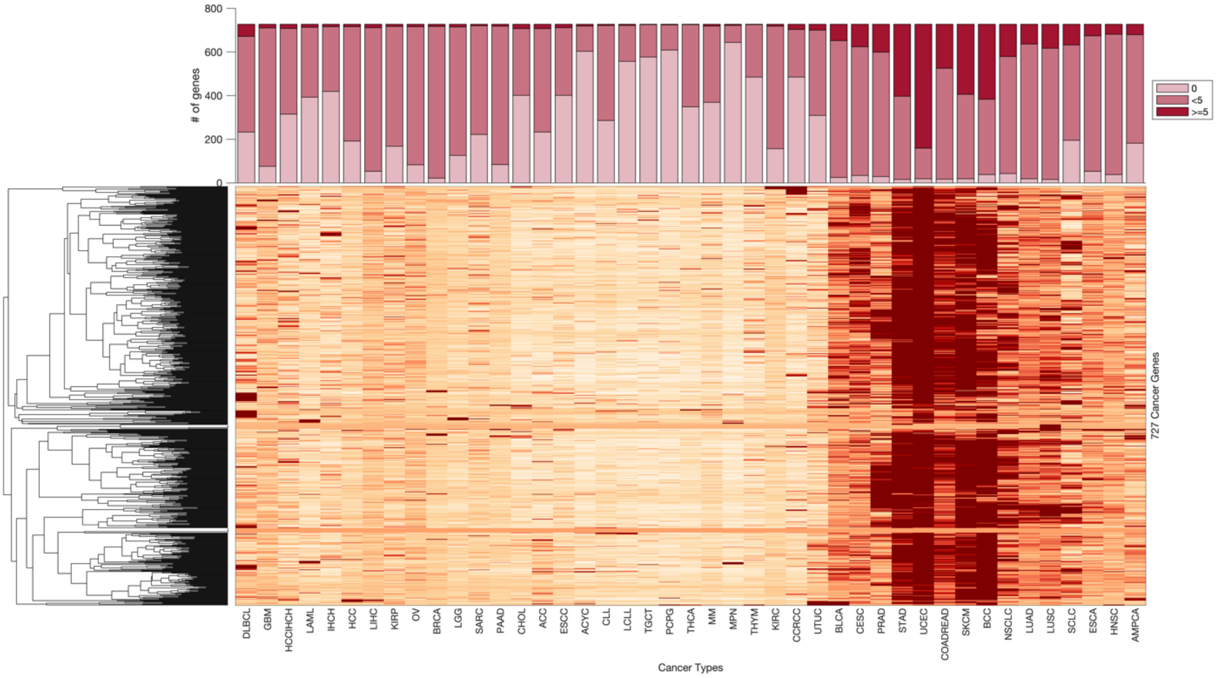


**Supplementary Figure 1:** *Unsupervised hierarchical clustergram of tumours based on the mutation frequency of cancer driver genes. The grouped bar graph at the top of the clustergram shows the number of cancer driver genes that are mutated in >= 5% of the samples for the darker coloured bar, in < 5% of the tumours for the intermediate shade), and none (0) for the lighter coloured bars.*

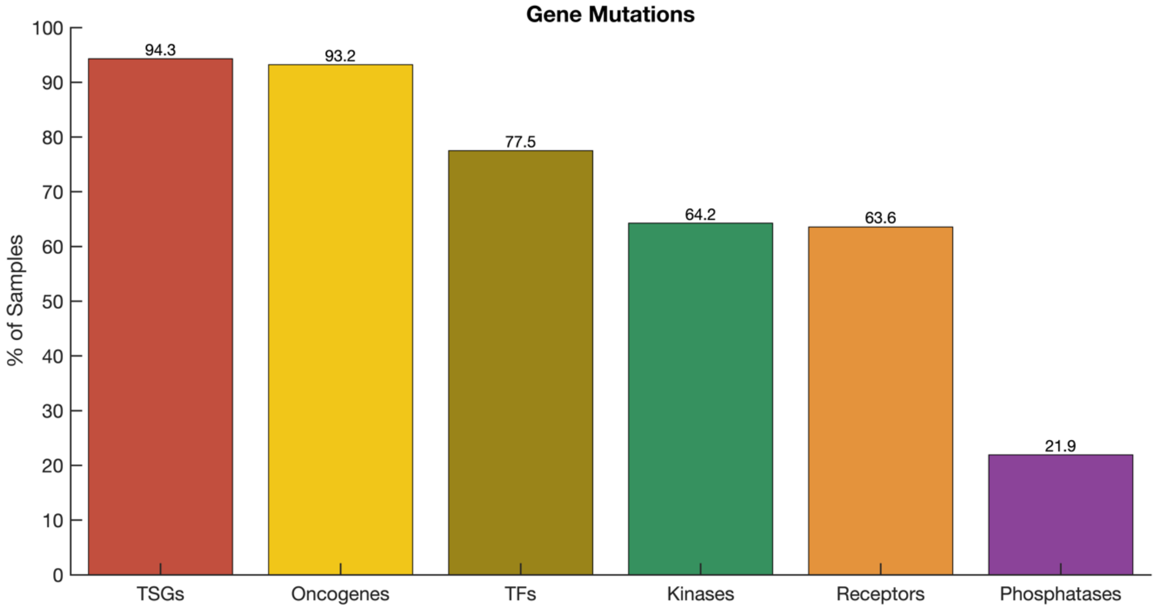


**Supplementary Figure 2:** *Frequency of driver cancer genes that belong to various classes across all tumour samples.*

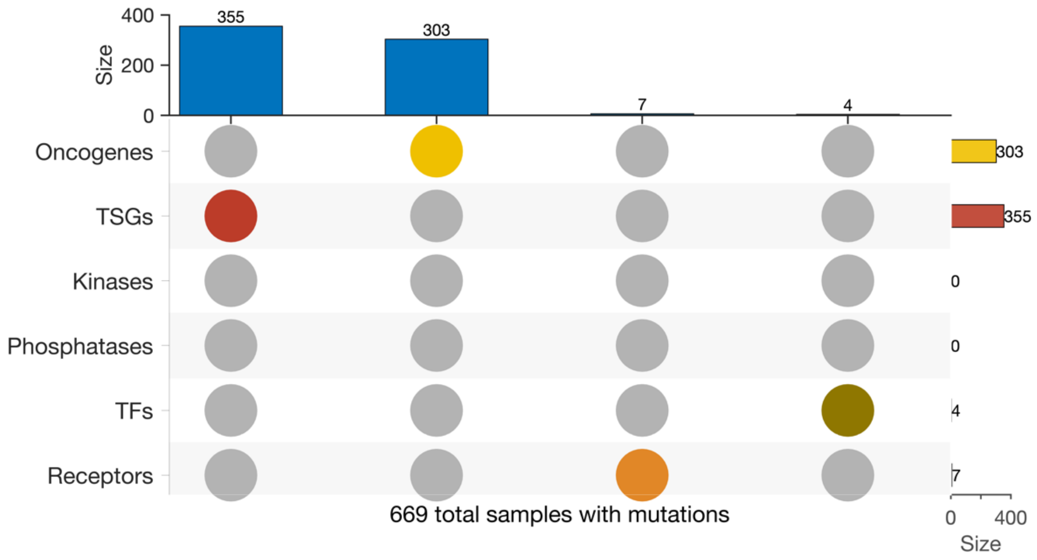


**Supplementary Figure 3:** *Upset plot showing the number of mutually exclusive mutated driver genes across all the tumour samples.*

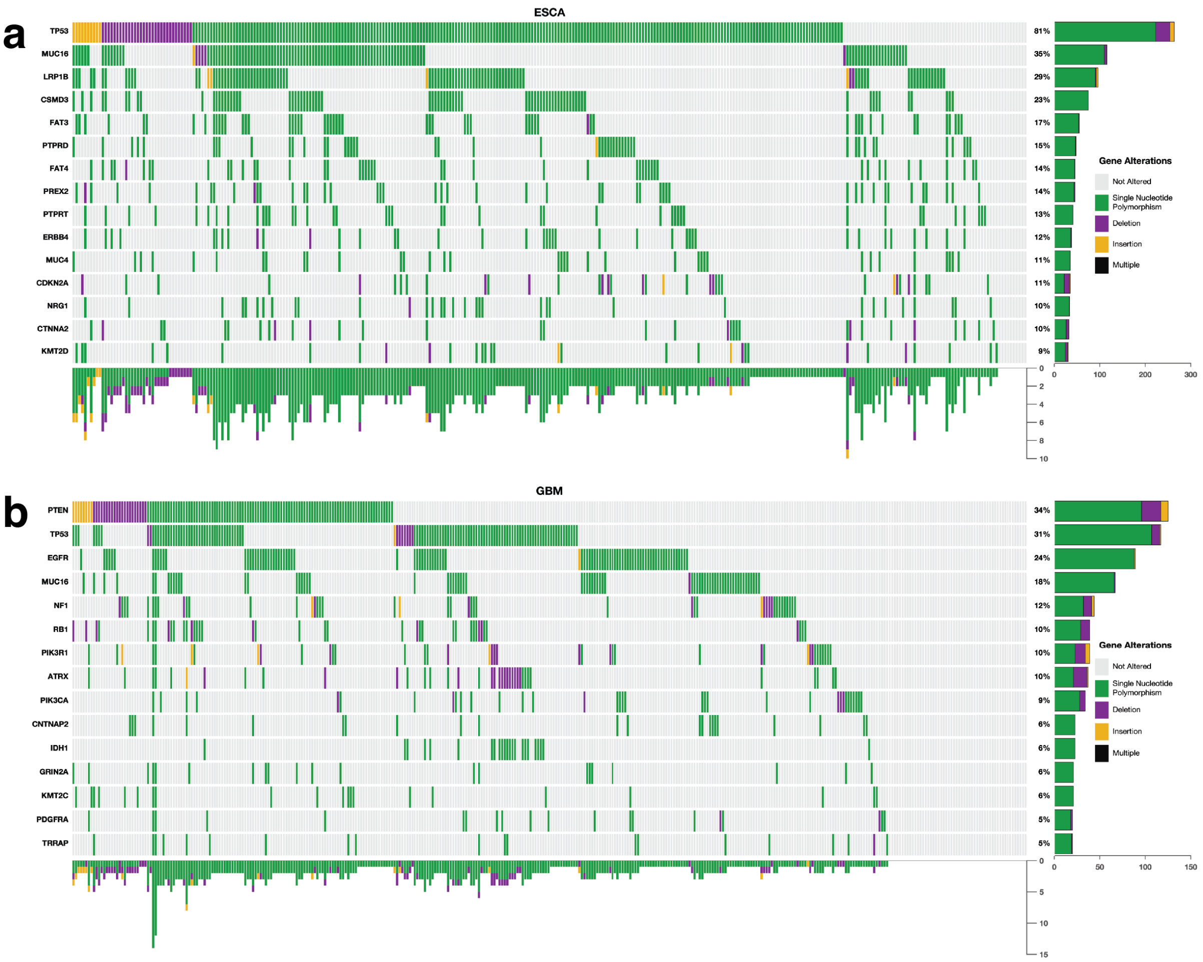


**Supplementary Figure 4:** *Example of mutation signature plot showing co-occurrence and exclusivity of mutations in (a) oesophageal cancer and (b) kidney cancer. The plot shows only the 15 most mutated genes.*

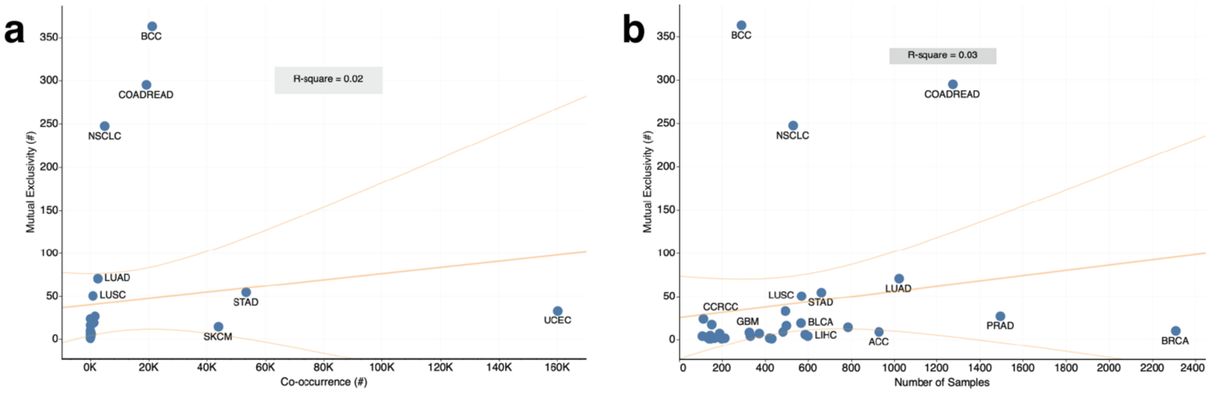


**Supplementary Figure 5:** *Correlation between the number of (a) co-occurring mutations and exclusive mutations in gene pairs observed and (d) exclusive mutations in gene pairs versus the number of samples profiled for each cancer type.*

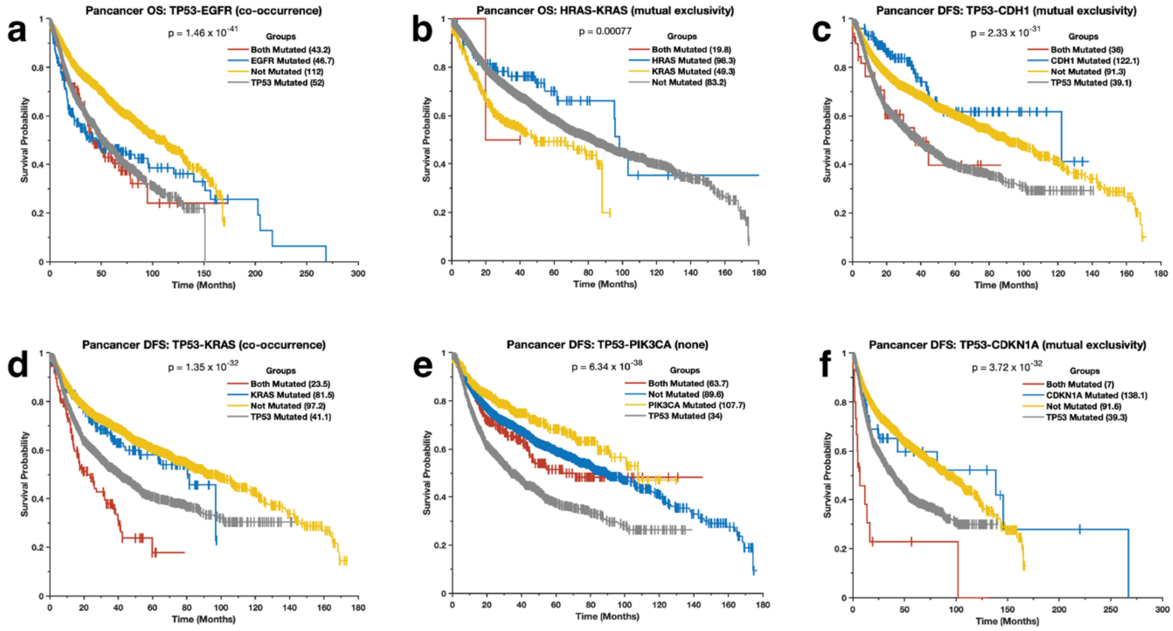


***Supplementary Figure 6:*** Kaplan-Meier curve of the disease-free and overall survival periods of patients afflicted by tumours with two cancer driver gene pairs mutated, only one gene of the pair mutated, and none of the genes in the pair mutated.

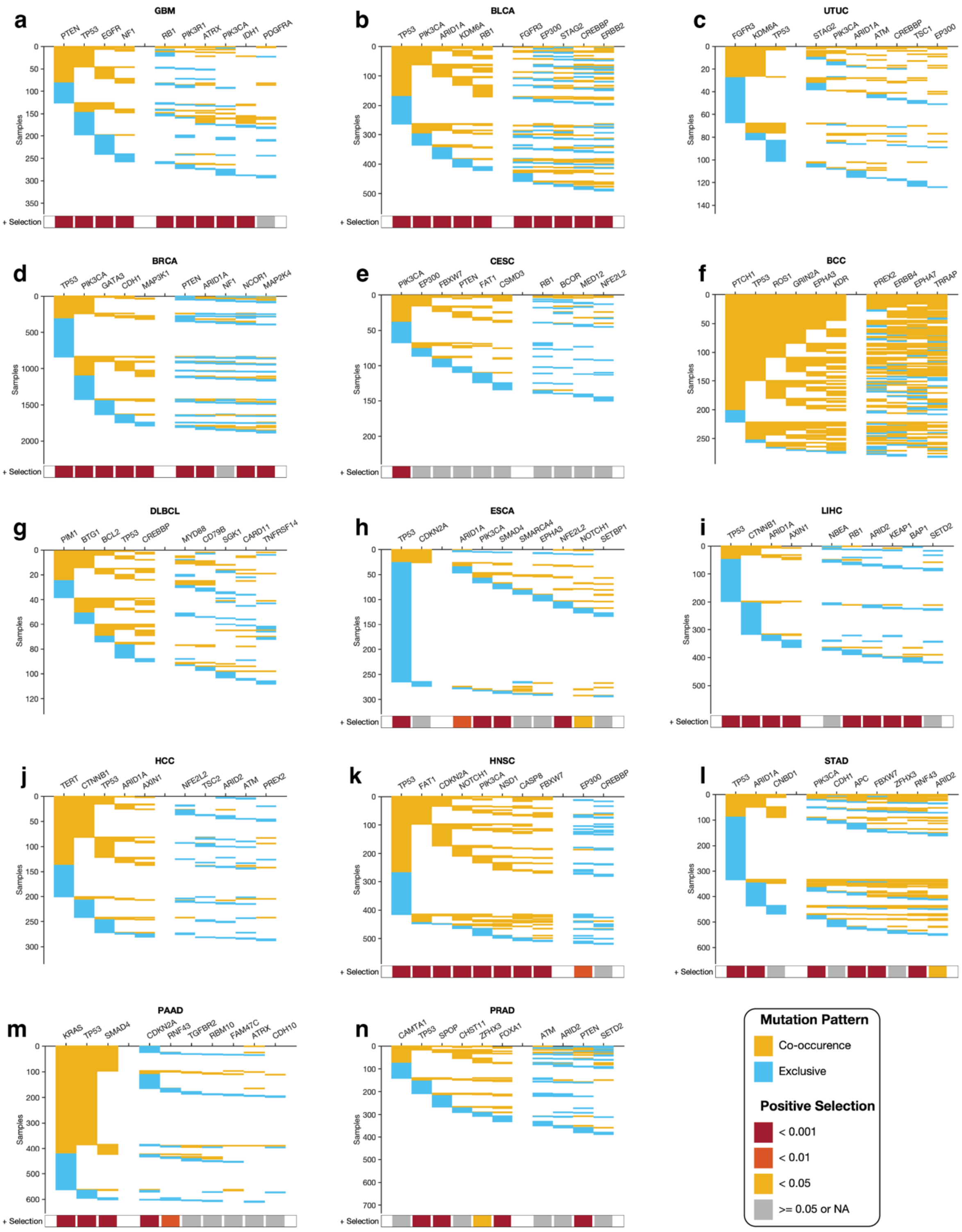


**Supplementary Figure 7:** *The two co-occurring pathways predicted to drive oncogenesis based on the mutational landscape in various cancer types. The coloured square marks at the bottom of each plot show a positive selection of mutations in each gene along each column (see Methods section). The connectivity of network components within each panel was extracted from the KEA and ChEA databases and the UCSC super pathway.*
